# Evidences for functional trans-acting eRNA-promoter R-loops at Alu sequences

**DOI:** 10.1101/2021.02.17.431596

**Authors:** Xue Bai, Feifei Li, Zhihua Zhang

## Abstract

Enhancers modulate gene expression by interacting with promoters. Models of enhancer-promoter interactions (EPIs) in the literature involve the activity of many components, including transcription factors and nucleic acid. However, the role that sequence similarity plays in EPIs, remains largely unexplored. Herein, we report that Alu-derived sequences dominate sequence similarity between enhancers and promoters. After rejecting the alternative DNA:DNA and DNA:RNA triplex models, we proposed that enhancer-associated RNAs, or eRNAs, may directly contact their targeted promoters by forming trans-acting R-loops at those Alu sequences. We showed how the characteristic distribution of functional genomic data, such as RNA-DNA proximate ligation reads, binding of transcription factors, and RNA-binding proteins, align with the Alu sequences of EPIs. We also showed that these aligned Alu sequences may be subject to the constraint of coevolution, further implying the functional significance of these R-loop hybrids. Finally, our results showed that eRNA and Alu elements associate in a manner previously unrecognized in the EPIs and the evolution of gene regulation networks in mammals.

## Introduction

The human genome consists of some 20000 protein-coding genes, while more than a million enhancers were estimated (Heintzman *et al*, 2009). Most phenotype-associated single nucleotide polymorphisms (SNPs) and genome structure variations mapped in the human genome were believed to be associated with enhancers (Ernst *et al*, 2011). Thus, the structure, origin and mechanism of enhancers are fundamental topics in modern molecular biology.

Several independent, but not mutually exclusive, models have been proposed to explain how enhancers interact with their targeted promoters. Promoter expression can be directly affected by physical contact with enhancers (Carter *et al*, 2002). These contacts may involve multiple transcription factors, such as mediator (Kagey *et al*, 2010), CTCF (Bell *et al*, 1999), and cohesion (Kagey *et al.*, 2010), participating in the formation of chromatin loops. The loop extrusion model has attracted much attention recently (Sanborn *et al*, 2015), as suggested by the observation that the binding of CTCF at loop anchors exhibited a specific convergent orientation (Guo *et al*, 2015; Rao *et al*, 2014). The loops were likely restricted within the structure of chromosomal domains, such as topologically associating domains (TAD) (Dixon *et al*, 2012; Nora *et al*, 2012), as defined by chromosome conformation capture-based technologies, e.g., Hi-C (Lieberman-Aiden *et al*, 2009) and ChIA-PET (Fullwood *et al*, 2009). Another model proposed the existence of a so-called transcription factory whereby RNA polymerase II (Pol II) could bind to both enhancers and promoters such that Pol II was unevenly distributed in nuclei, forming discrete nuclear foci (Sutherland & Bickmore, 2009). The clustering of Pol II would pull enhancers and promoters into these transcription factories where they would make contact. In addition to direct physical enhancer-promoter interactions, promoter activity may also be regulated indirectly by enhancers through RNAs associated with enhancers known as eRNAs (Kim *et al*, 2010). In one plausible model, eRNAs act as a scaffold to mediate the formation of a complex of EPIs (Hsieh *et al*, 2014). A related, but the much more sophisticated liquid-liquid phase separation model has recently emerged in the literature (Hnisz *et al*, 2017). Here, liquid condensates could be formed by components, such as the C-terminal domain of Pol II (Boehning *et al*, 2018), transcription factors (TFs), and coactivators that bind on both enhancers and promoters (Boija *et al*, 2018). However, some fundamental questions remain unanswered. For example, in addition to binding with transcription factors, the potential role of DNA/RNA sequences in EPI functionality is worthy of investigation.

In mammals, a large portion of the genome was derived from transposable elements (TE), about 45% in the human genome. The L1 and Alu elements are the most abundant mobile elements comprising 21% and 11% of the human genome, respectively (Batzer & Deininger, 2002). Based on their potential binding by TFs (Cordaux & Batzer, 2009), TEs have long been regarded as a major source of innovation in gene regulatory networks (Feschotte, 2008). For example, about 25% of human promoters were found to contain TE-derived sequences (Jordan *et al*, 2003), implying a possible origin from TE. Accumulated evidences in the literature supports TE origination of enhancers, as well (Feschotte, 2008). For example, since Alu elements were found in most enhancers, they were suggested as proto-enhancers in primates (Su *et al*, 2014). The association of Alu elements with both enhancers and promoters implied sequence similarity among cis-regulators. However, the functional indications of this sequence similarity remain unexplored.

In the present work, we reported that Alu elements dominated the aligned sequences between enhancers and promoters. Further evidences suggested that the concentration of Alu-derived sequences in both enhancers and promoters may be subject to stable selection during evolution. After excluding alternative explanations, e.g., DNA-DNA direct hybrids and DNA-RNA triplex, we showed evidences to support a trans-acting R-loop model. This model proposes the existence of functional eRNA-promoter R-loop hybrids in EPI pairs with identical Alu-derived sequence fragments. These results, in turn, suggested a new role for eRNAs in the regulation of enhancer-promoter interactions. Finally, our results showed that eRNA and Alu elements associate in a manner previously unrecognized in EPI regulation and the evolution of gene regulation networks in mammals.

## Results

To identify enhancers, their eRNAs, and potential targeted promoters, we composed a simple pipeline to retrieve genome loci with characteristic features in transcriptome and epigenome data (see Methods, Fig EV1A). Briefly, active enhancers were defined as genome regions that overlapped with peaks of DNase I hypersensitive sites (DHSs), ChIP-seq peaks of histone modification (H3K27ac and H3K4me1), and peaks of global nuclear Run-On sequencing (GRO-seq). Potential transcription starts sites (TSS) of eRNAs were defined by cap analysis of gene expression and deep sequencing (CAGE-seq) data. Promoters were defined as the 3kb flanking region of annotated TSS (TSS-2KB, TSS+1KB). All intra-TAD enhancer-promoter pairs were assigned as potential enhancer-promoter interactions (EPI, Fig EV1B). In total, we identified 3858 active enhancers with the average length 1.018kb, 6839 active promoters and 18807 enhancer-promoter interactions (EPI) in GM12878 cells using the data from Encyclopedia of DNA Elements (ENCODE) Project (Consortium, 2012). In support of the accuracy of this pipeline, most (97.3%) of the predicted enhancers were annotated as strong enhancers by ChromHMM(Ernst & Kellis, 2017), and the enrichment of ChIA-PET loops and Hi-C reads supported the predicted EPI compared to the randomly rewired controls (Fig EV1E, Fig EV1F).

### Alu elements are strongly associated with enhancer-promoter interactions

To investigate the sequence-related association between enhancers and promoters in EPIs, we BLASTed the paired enhancers and promoters. About 31.6% of enhancers and 42.4% of promoters were found to carry at least one BLAST best hits (E-value < 1E-4). Intriguingly, almost all the BLAST hits (98.70%) were part of the Alu elements (Fig 1A), resulting in a remarkably high correlation between the distribution of BLAST hits and Alu elements in both enhancer and promoter regions (Pearson correlation coefficient, PCC = 0.969 and 0.987, respectively. Fig 1B). Further MEME analysis of the BLAST hits showed a group of enriched sequence motifs, and two out of the top five motifs (motif1 and 2) were known to be conserved in Alu elements (Kariya *et al*, 1987) (Fig 1C-1D, Table EV1). Making comparisons with the JASPER database (Fornes *et al*, 2020), we found that all five enriched sequence motifs matched well with known TF-binding motifs, e.g., GFi1b, SREBF2 and ZNF263. Therefore, we asked if this overwhelmingly dominant Alu sequence association between enhancers and promoters could be biologically relevant. Biological relevance supposes a distinct distribution of Alu elements in the enhancers and promoters from genomic background. Surprisingly, when we compared their flanking genome context, we found that the Alu elements were infrequently settled in both the promoters and enhancers. Average Alu concentration in the flanking genome regions was 1.64- and 1.68-fold higher than that in the enhancers and promoters, respectively (Fig 1E). The concentration in the enhancers was even 1.2-fold lower than the whole genome average (Fig 1E). Interestingly, Alu elements were not totally depleted from those cis-regulatory elements, but largely restricted to less than two (Fig 1F). By percentage, 38.77% and 43.44% of enhancers and promoters carry one or two Alu elements, while only 1.35% and 26.64% carry more than two Alu elements, respectively (Fig 1F. Appendix Table S1-S4). This partial depletion of Alu elements is also true for the super-enhancers (SE) (Fig 1F); that is, compared to 22.91% in the SE flanking region, 26.02% (427 out of 1641) of the SE containing normal enhancers carry exactly one Alu (P=2e-4, χ^2^ test). Thus, the depletion of Alu elements hints that the number of Alu elements may be subject to some evolutionary constraint within those cis-regulatory regions. Given this observation, we speculated that the Alu elements may be associated with regulatory EPIs.

**Figure 1.**
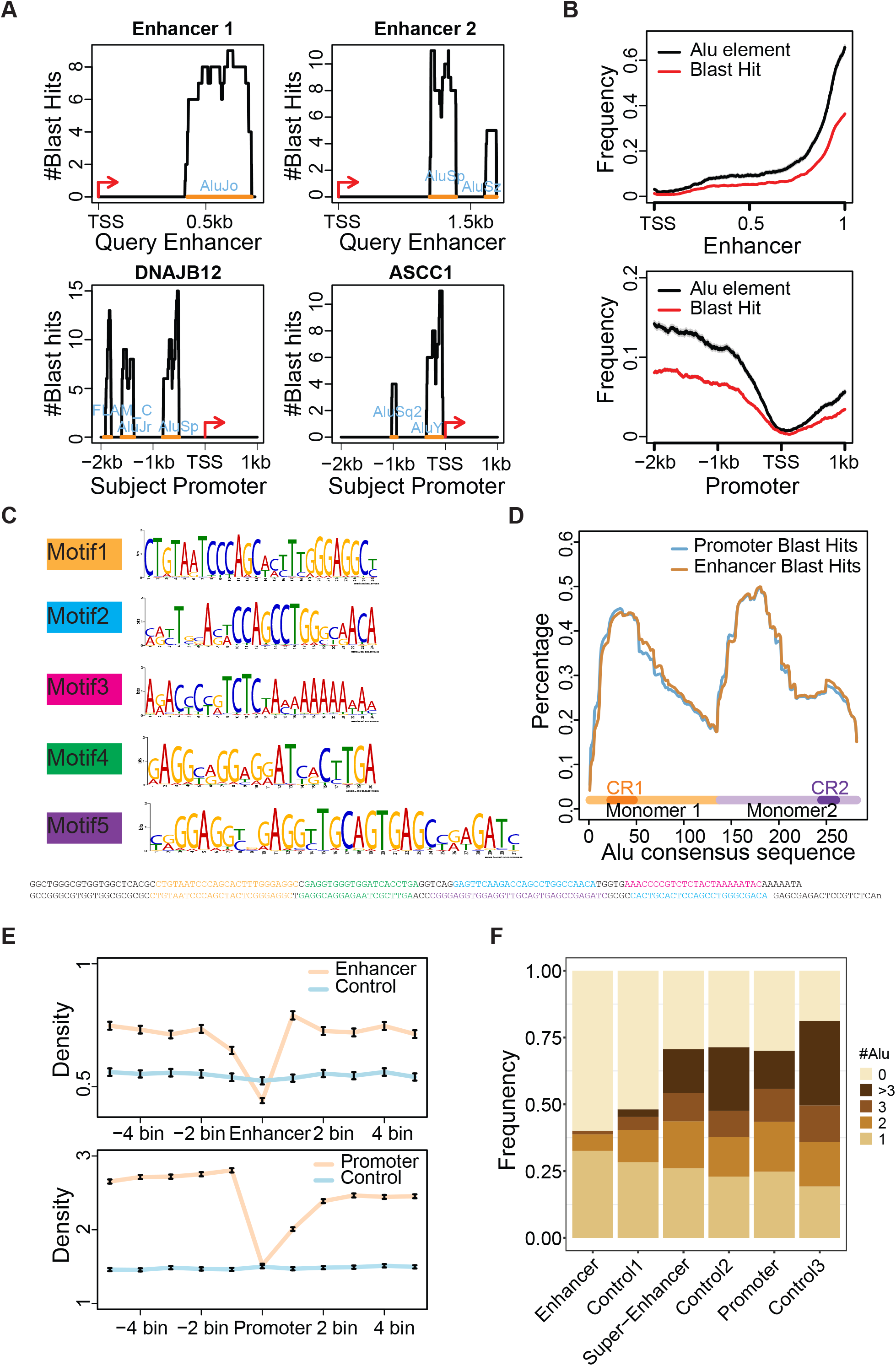
Alu-derived elements dominate enhancer-promoter alignment. **A.** Examples of enhancer-promoter Blast alignment. TSS in both enhancers and promoters were defined by CAGE data. **B.** Distribution of Blast hits between enhancer-promoter and Alu-derived elements in promoters and enhancers. The enhancers were rescaled uniformly. **C.** The five enriched sequence motifs found in the Blast hits between the enhancers and promoters. Motif1 and motif2 are enriched in the conserved regions (CR) of Alu elements; Motif3 is similar to the binding motif of DNA-binding protein ZNF384, and Motif5 is enriched in the insertion region of second monomer of the Alu elements. **D.** Distribution of Blast hits on the Alu consensus sequence. CR1 and CR2 refer to the first and second conserved region (23-47 and 245-260), respectively. **E.** Distribution of Alu-derived sequences in the enhancers and promoters. Each bin represents a genome fragment with the length of the given promoter or enhancer. The control was randomly sampled genome fragments with length identical to that of promoters and enhancers. The error bars represent STD/SQRT(n). **F.** Distribution of the number of Alus in the enhancers, super-enhancers and promoters. Control1, control2 and control3 are the randomly sampled genome fragments with distribution of number and length identical to that of enhancers, super-enhancers and promoters, respectively.

### Alu elements may be involved in the regulation of gene expression

If Alu elements are involved in the regulation of EPIs, then distinct expression levels between genes with and without Alu association in their cis-regulators can be expected. To test this speculation, we classified EPIs into four groups according to whether the enhancers (E) and promoters (P) carry Alu elements (+) or not (−), and we compared the expression pattern of genes among the groups.

First, we found that the expression level of genes with an Alu element in their promoter (denoted as E*P+I) was significantly higher than that of genes not carrying an Alu element in their promoter (denoted as E*P-I, Mann-Whitney test; p=3e-10, Fig 2A). Within the E*P+I genes, the expression of genes with an Alu in their enhancers was also higher than that that of genes without an Alu element in their enhancer (denoted as E+P+I *vs*. E-P+I, Mann-Whitney test; p=0.030). Meanwhile, little, to no, effect of Alu elements on enhancers was observed for the expression level of genes without an Alu element in the promoters (denoted as E+P-I *vs*. E-P-I, Mann-Whitney test; p=0.066 Fig 2A). The high expression level for E-P+I genes could be explained by the presence of alternative enhancers, located in the same TAD and carrying an Alu, regulating the promoter, since a promoter can be regulated by multiple enhancers. Indeed, for E-P+I promoters, 43.9% have at least one such alternative Alu-carrying enhancer. This possibility was further evidenced by the indistinguishable expression level between E-P+I and E*P-I genes after removing the genes with multiple Alu-carrying enhancers from E-P+I (Mann-Whitney test, P-value=0.2, Fig 2A). This higher gene expression level in the E+P+I pattern can also be seen in K562, HepG2, HeLa-s3 and MCF-7 cell lines, according to ENCODE transcriptome data(Consortium, 2012) (Fig EV2A–EV2D).

**Figure 2.**
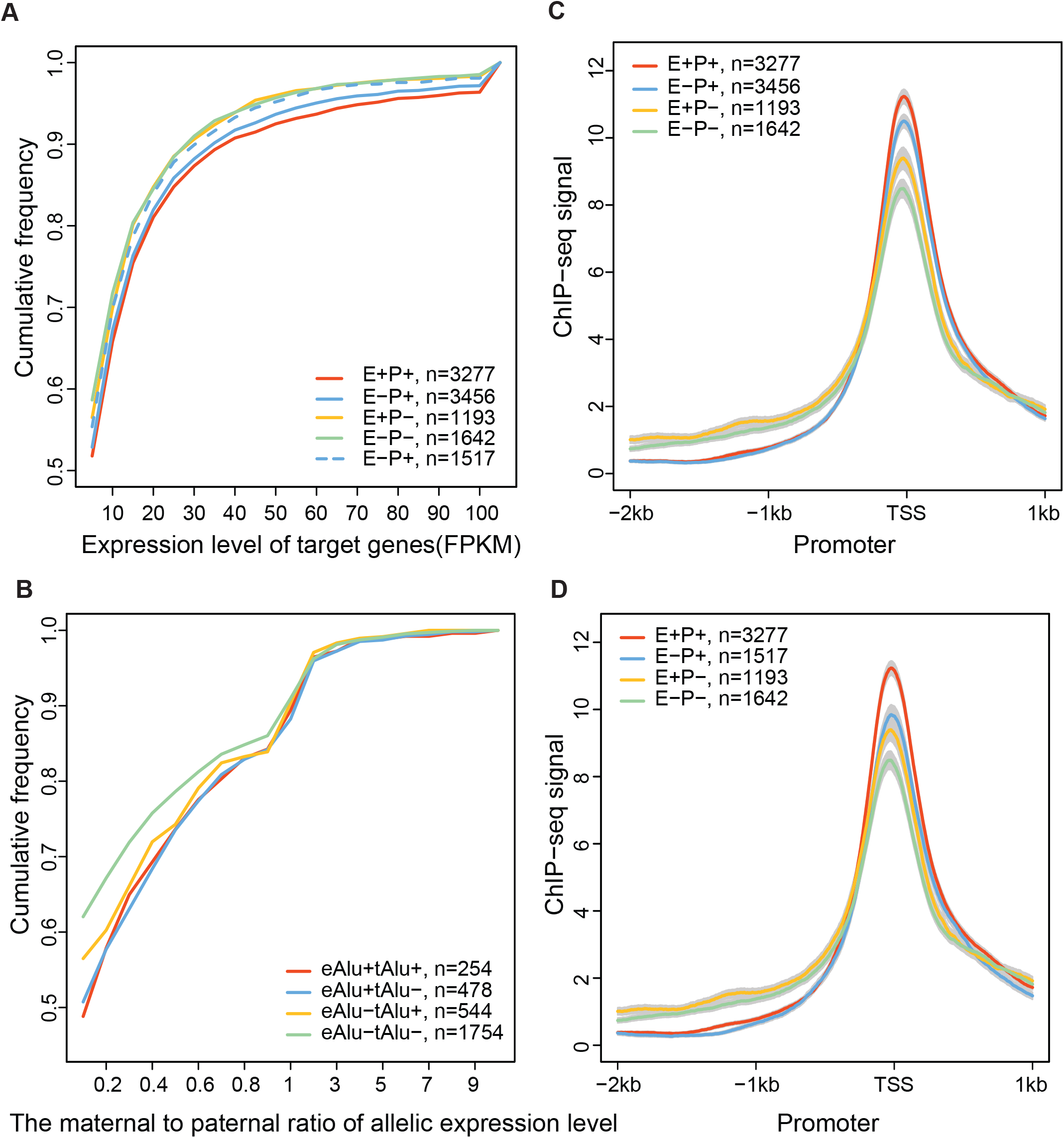
Distribution of gene expression level in different EPI categories. **A.** Cumulative frequency of gene expression in GM12878. E+ and E-represent enhancers that carry at least one or zero Alu element, respectively. P+ and P-represent the putative target promoters that carry at least one or zero Alu element, respectively. The blue dotted line represents the EPIs in the E-P+ group after removing the genes with multiple Alu-carrying enhancers. **B.** The cumulative frequency of differential expression between homologous alleles in GM12878. The maternal to paternal ratio of allelic expression level was calculated by the number of RNA-seq reads that can be assigned to maternal and paternal alleles by the SNPs. eAlu+ and eAlu-represent the enhancers carrying Alu with at least one or zero SNP, respectively. tAlu+ and tAlu-represent the putative target promoters carrying Alu with at least one or zero SNP, respectively. **C** and **D**. Distribution of TF ChIP-seq signals in the putative target promoters. The genes that have multiple enhancers were removed in **D**.

Second, cis-regulators with SNPs on their carried Alus (cAlu) are frequently involved in allelic differential gene expression. If cAlus are essential for gene regulation and if DNA sequence matching between cAlus in enhancers and promoters is critical, then we could expect two homologous alleles to be differentially expressed in the presence of heterozygous SNPs in their cAlus. To test this prediction, we compared the difference in allelic expression between genes with and without SNPs in their cAlus in GM12878 cells, which have 2 million well-phased SNPs. Indeed, irrespective of whether the SNP appears in promoter (pAlu+), enhancer(eAlu+) or both, the difference in expression level is always higher than that in genes without such SNPs, as long as a heterozygous SNP is present in the cAlu, (Mann-Whitney test, P-value=2e-08, Fig 2B).

Finally, more transcription factor (TF) bindings are found in Alu elements carrying promoters than those not carrying promoters. By integrating ChIP-seq data from the ENCODE project, we compared the ChIP-seq signal density of 86 TFs in the promoters among the four EPI groups (Fig 2C and 2D, Dataset EV2), and we found that the density in E*P+I promoters was higher than that in the E*P-I groups (Fold change = 0.21, Mann-Whitney test, P-value = 1.26e-21, Fig 2C). Again, when we removed the genes with alternative cAlu-carrying enhancers from E-P+I, the significance became substantially weaker (Fig 2D, Fold change = 0.18, Mann-Whitney test, P-value = 3.46e-16). Collectively, the above evidences supported our speculation that cAlus may be associated with the regulation of gene expression.

### Alu-mediated EPI disfavors direct DNA-DNA contact at the Alu element

We asked whether those cis-regulated DNA regions could directly contact each other at the cAlus. Direct DNA-DNA contact at cAlus would imply symmetric distribution of ChIA-PET loop anchors flanking the cAlus since the up- and downstream genome cAlu fragments would have an equal chance for ligation while preparing the ChIA-PET library. However, for both Pol II and CTCF data we examined, the ChIA-PET loop peaks were clearly skewed and asymmetrically distributed flanking the cAlus in both enhancers and promoters (Fig 3A). Thus, our data suggested that cAlu-cAlu direct DNA-DNA contacts may be disfavored, at least near the Pol II and CTCF ChIA-PET loop anchors. We then asked if this asymmetric distribution could have resulted from chromatin loop extrusion characterized by convergent binding of the CCCTC-binding factor (CTCF) at the chromatin loop anchors(Rao *et al.*, 2014). To test this speculation, we calculated the enrichment of CTCF motif orientation combinations between cAlu pairs. However, few signs gave evidence of the enrichment of converging CTCF motifs. In fact, we saw a slight depletion from the paired cAlus (odds=0.87, Fig 3B). Moreover, none of these convergent motif pairs overlapped with CTCF ChIP-seq peaks or ChIA-PET loop anchors. To exclude the possible false peak calling owing to low mappability at the Alu element, we recalled the ChIP-seq peaks included in multi-mapped reads (see Methods). However, very few ChIP-seq peak-pairs at convergent CTCF motif pairs were found with those multi-mapped reads. Thus, loop extrusion is less likely to be a dominant mechanism underlying cAlu-cAlu association. Taken together, direct cAlu-cAlu DNA contacts may be disfavored between the cAlu-carrying cis-regulators.

**Figure 3.**
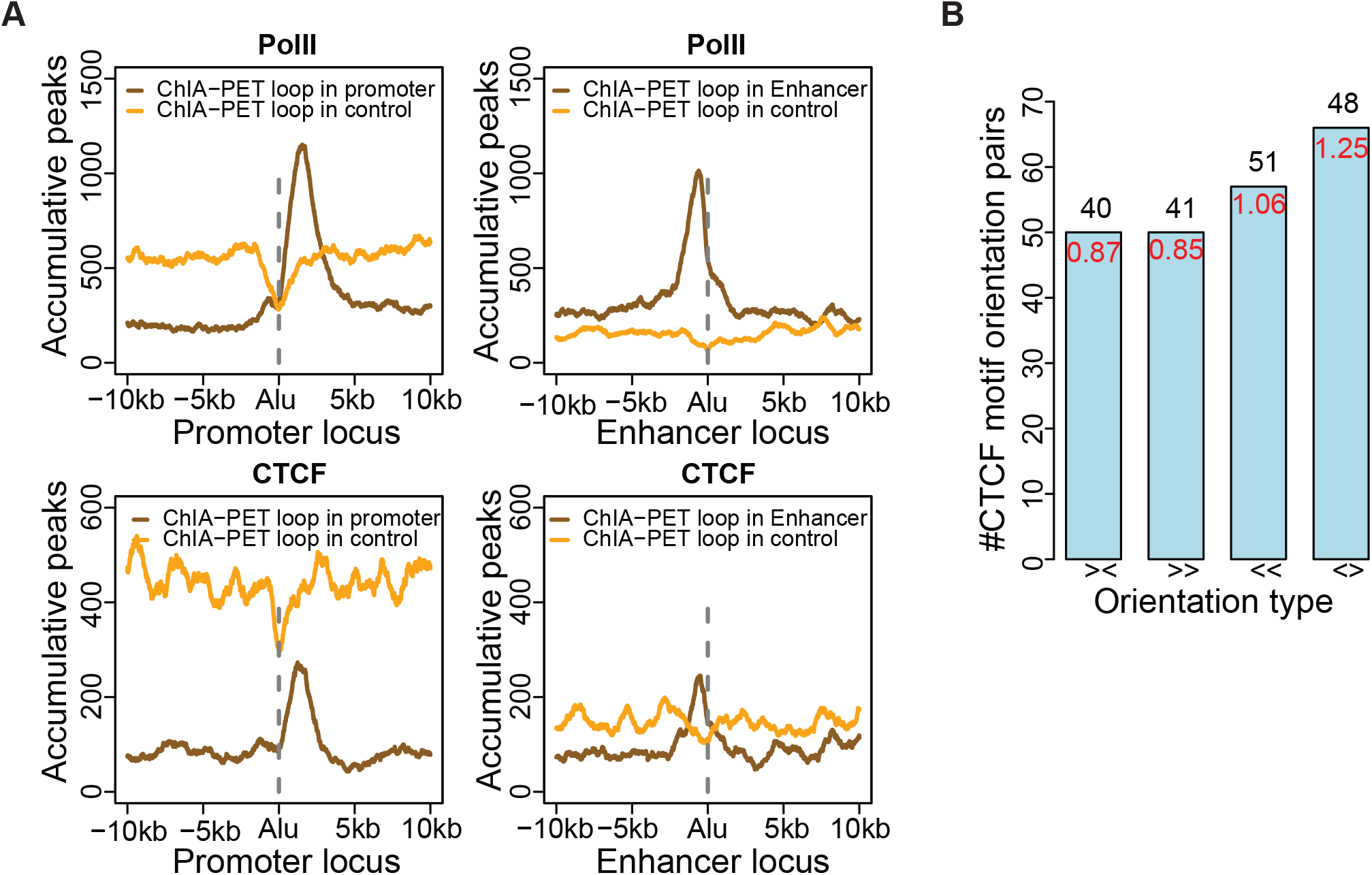
Distribution of ChIA-PET loop and CTCF orientation around cAlus. **A.** Distribution of ChIA-PET peaks flanking the cAlus. Orientation of enhancers was defined by CAGE, and the bidirectional enhancer was treated as two enhancers. **B.** Number of CTCF motif pairs in cAlus. The black number on top of each bar indicates the number of EPIs. The enrichment of CTCF motif orientation pairs at cAlu-cAlu over the whole EPI (see Methods).

### Alu elements may mediate interaction between eRNAs and promoter through RNA-DNA

Next, we asked if cAlus interact *in-trans*, i.e., make contact through eRNAs. We found that cAlus do interact *in-trans* for two reasons. First, here, we support this speculation in two ways. First, RNA fragments that can be approximately ligated to DNA *in vivo* are prone to flanking the Alu elements. If the eRNAs are, indeed, hybrids to the promoters at the Alu elements, we would expect the sequencing reads from spatial approximation ligation-based technologies, e.g., GRID-seq(Li *et al*, 2017) and iMARGI(Yan *et al*, 2019), to concentrate flanking at the cAlu elements (Fig 4A). In the GRID-seq data from MDA231 and MM1s cells(Li *et al.*, 2017), 997 EPIs had at least one pair of GRID-seq reads supported. Among those 997 EPIs, we compared the distribution of reads flanking the cAlus and their genomic background. A clear bimodal distribution flanking the cAlu of the enhancer regions was observed (of 997 EPIs, P-value<2e-16; two-tailed chi-squared test (Fig 4B top)). The depletion of GRID-seq reads at cAlu was not caused by the poor mappability of the Alu loci (Fig 4B bottom), as, on average, only 1 pair of multi-mapped reads was involved in the enhancers we investigated. The reads from DNA end of GRID-seq were not included in this analysis, because they were generated by the AluI restriction endonuclease, which has preference to cut the Alu element.

**Figure 4.**
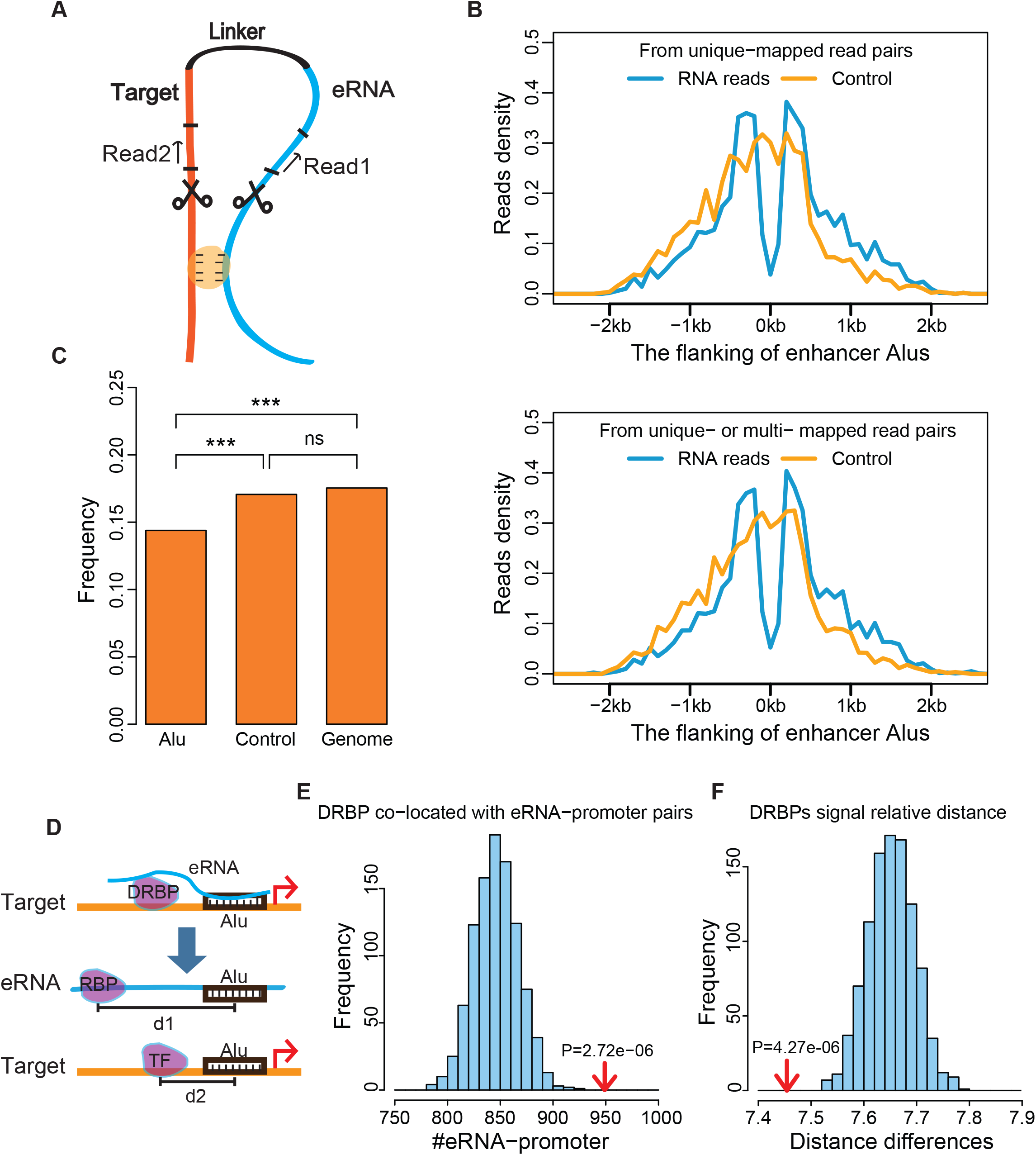
Distribution of GRID-seq and iMARGI reads flanking cAlu elements. **A.** Schematic diagram of sequencing technology for RNA and DNA interaction. The yellow oval represents interaction sites, which may or may not be mediated by RNA:DNA hybrids. **B.** Distribution of GRID-seq RNA reads (read1) around eRNA Alu elements. The top and bottom panels were plotted with unique mapped and total reads, respectively. **C.** Frequency of the iMARGI DNA read (read2) in the cAlus of putative target promoters and controls. **D.** Schematic diagram of TF- and RBP-binding model in the Alu-mediated eRNA:Promoter contacts. The d1 represents the genome distance between the middle of RBP eCLIP peak and Alu middle position in eRNA; d2 represents the genome distance between middle of TF ChIP-seq peak and Alu middle position in promoter. **E.** Distribution of number of EPIs with DRBP-binding on both eRNA and promoter side. The red arrow indicates the observed number (z-score=4.54), while the bars represent the distribution of 1000 randomly rewired E-P pairs with DRBP-binding on both eRNA and promoter. The CLIP-seq peak has to overlap with the extended region of enhancer transcript (+/−500bp), and the ChIP-seq peak has to overlap with the extended region of promoter region (+/−500bp). **F.** Distribution of distance difference between d1 and d2. The red arrow indicates the observed difference (z-score=−4.45), while the bars represent the distribution of the distance differences from 1000 randomly rewired E-P pairs. The eCLIP-seq peaks have to be overlapped with the extended region of enhancer transcript (+/−500bp), while the ChIP-seq peak has to be overlapped with the extended region of promoter region (+/−10kb), followed by choosing the peak nearest cAlu. ***:P-value<0.001; U test.

To perform this test with iMARGI data, we aligned all the distal (>2000bp) unique mapped DNA reads to the promoters with and without cAlus (Methods), and we found a significant depletion in Alu elements in the promoters with cAlus (P-value<2e-16, two-tailed χ^2^ test, Fig 4C). Thus, both GRID-seq and iMARGI data suggested that the Alu elements may settle in the middle of RNA:DNA-interacting loci.

RNA-DNA interacting at the cAlu loci was further evidenced by the RNA-protein interaction assay. One type of TF can bind to both DNA and RNA, named DNA and RNA binding proteins (DRBPs). If cAlu-cAlu contacts mediate EPI, we would expect DRBP-binding to be prone to EPI sites (Fig 4D). To test this speculation, we collected 22 DRBPs with both ChIP-seq and eCLIP-seq data available in K562 cells from ENCODE(Consortium, 2012) (Dataset EV3). We saw significantly enriched EPIs with eCLIP-seq and ChIP-seq peaks located in their enhancer and promoter loci, respectively and simultaneously, compared to those in the randomly assigned promoters with similar expression level (Fig 4E). Further, we would expect DRBPs to bind at, or flank, the cAlu-cAlu contacting loci, resulting in similar binding sites to the Alu distances between the RNA and DNA (Fig 4D). By comparing the distance of DRBP-binding sites to the cAlus in eRNA and promoters, we found that the differences in distance were significantly less than those of random control (P-value<0.001, Fig 4F). Together, these data suggest that RNA:DNA interaction might occur through Alu:Alu pairing in the EPI. Two types of RNA-DNA interactions are DNA:RNA triplex (Felsenfeld & Rich, 1957) and R-loops (Thomas *et al*, 1976). We next examined which type of RNA-DNA interaction is more plausible in Alu-Alu-mediated EPI.

### The DNA:RNA triplex may not be favored at the cAlus

DNA:RNA triplex (DRT) may have an important regulatory function, e.g., RNA-mediated target-site recognition (Senturk Cetin *et al*, 2019). To examine if DRT is prevalent in Alu-mediated EPI, we predicted 3,164,635 potential DRTs at the predicted EPI loci using Triplexator (Buske *et al*, 2012). However, we found that those predicted DRTs were depleted from cAlus. The predicted DRTs were almost evenly distributed in eRNA and promoter loci and only weakly enriched in the upstream of promoter TSS (Fig 5A). Compared with the flanking region of cAlus, the number of predicted DRTs were significantly lower at the cAlus in both eRNA and promoter regions (P-value<2e-16, Mann-Whitney test, Fig 5B).

**Figure 5.**
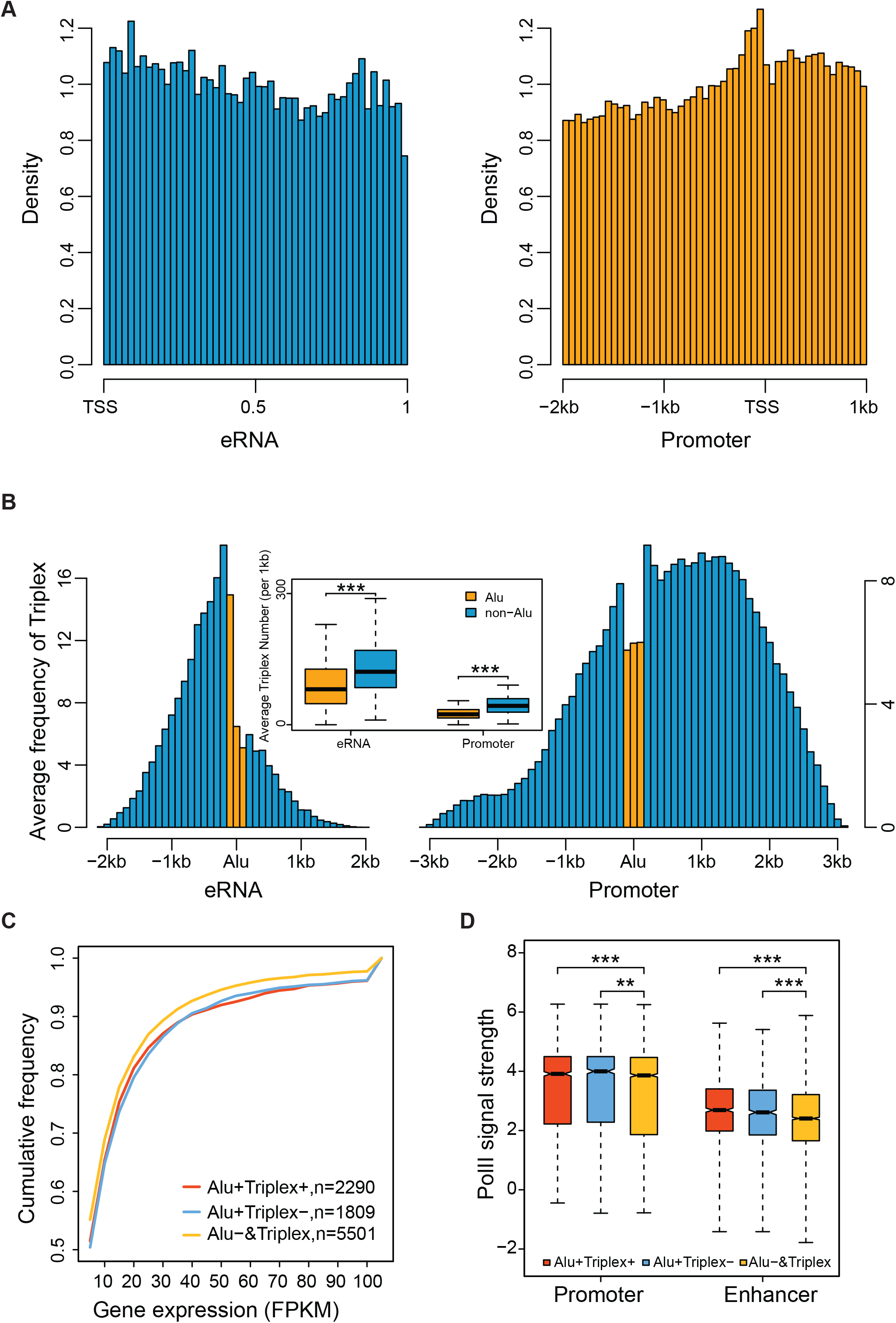
Distribution of predicted interaction of RNA:DNA triplex in EPI. Distribution of predicted RNA:DNA triplex in the eRNAs and putative target promoters. The RNA:DNA triplex was predicted by Triplexator with minimal length 10bp. **A.** The eRNAs were aligned and rescaled uniformly, and promoters were aligned. **B.** The eRNAs and promoters were aligned centered at the Alu elements, which were marked in orange. The average number of predicted RNA:DNA triplexes in Alu and flanking regions were shown in the embedded plot. **C.** Cumulative distribution of gene expression level for the putative target genes. Triplex+ and Triplex-represent with and without triplex predicted at the cAlus, respectively. &Triplex represents with triplex predicted. Alu+ and Alu-represent with and without cAlus, respectively. (P-value=4e-05 and 1e-06, respectively) **D.** The distribution of Pol II ChIP-seq signal strength in the transcriptional enhancers and putative target promoters. (P-value= 6e-04 and 9e-03, respectively) **, P-value <0.01; ***, P-value < 0.001, Mann-Whitney test.

Next, we examined the association between DRT-carrying Alus and gene expression. We divided DRTs involved in EPIs into three groups: EPIs carrying Alus that also carry a DRT (Alu+Triple+, n=3757), EPIs carrying Alus that carry no DRT (Alu+Triplex-, n=2526) and EPIs carrying no Alu, but have DRT (Alu-&Triplex, n=12524). By comparing the expression level of the genes among the groups, we found that the expression level and the binding of Pol II in the groups carrying Alu (Alu+Triplex*) were significantly higher when compared to the group carrying no Alu (Alu-&Triplex, Mann-Whitney test; P-value <0.01). However, little difference was seen between the groups having a DRT, or not, in the Alu (Fig 5C-5D). Based on these results, our analysis can reject the possibility that DRT plays a role as a main mechanism in Alu-mediated EPI.

### R-loop concentrated at the Alu element

We next examined whether Alu-mediated EPI occurs through RNA-DNA hybrids, which is also known as R-loops. First, we defined a set of reliable trans-R-loops from strand information based on single-strand DNA ligation-based library construction from DNA:RNA hybrid immunoprecipitation, followed by sequencing (ssDRIP-seq (Xu *et al*, 2017)) data for HeLa cells (Yang *et al*, 2019) (Fig 6A). Briefly, the cis- and trans-R-loops were defined as follows. The ssDRIP-seq pair-end reads were defined as cis if read2 was mapped to the sense strand of the locus where read1 was mapped. The ssDRIP-seq pair-end reads were defined as trans if read2 was mapped to the anti-sense strand of the locus where read1 was mapped and no transcription activity was detected in the anti-sense strand, as indicated by RNA-seq (Consortium, 2012) and GRO-seq data (Bouvy-Liivrand *et al*, 2017) (Fig 6A). The overlapping genes were not included in this analysis. For the three replications of ssDRIP-seq data in HeLa cells, we identified, on average, 315,167 and 395,532 trans and cis ssDRIP-seq reads, respectively, from 9,313 promoters of protein-coding genes.

**Figure 6.**
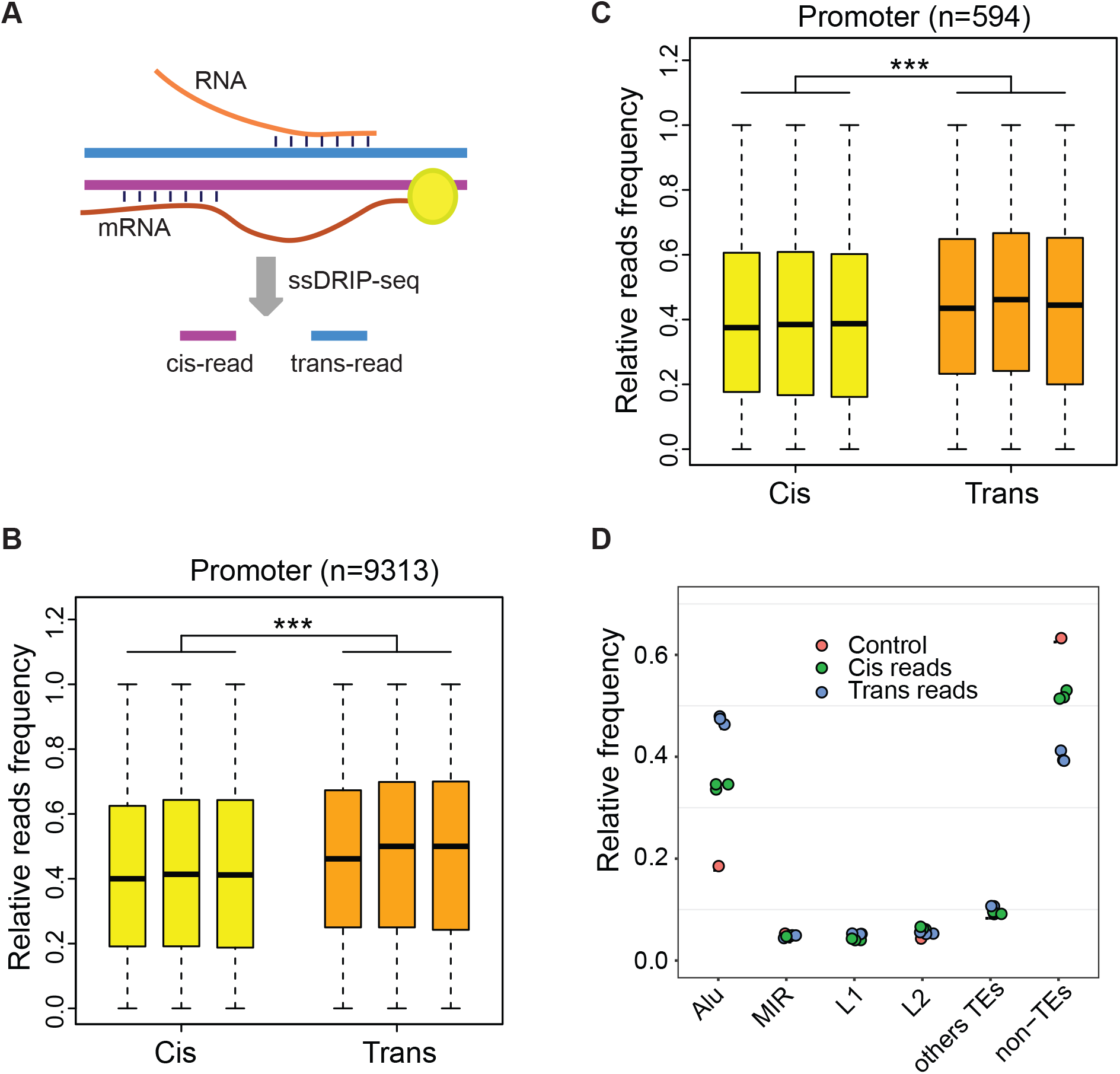
Distribution of *trans* ssDRIP-seq reads in promoters. **A**. The definition of *cis* and *trans* ssDRIP-seq reads. The *cis*- and *trans-*reads are those reads mapped to the template and sense strand of mRNA, respectively. For *trans*-reads, we further asked if the sense strand has no detectable transcript activity. The *trans* ssDRIP-seq reads are significantly enriched, overlapping cAlus in **B** promoter and **C** putative target promoter, compared to *cis* ssDRIP-seq reads in HeLa. **D**. Percentage of the *trans* and *cis* ssDRIP-seq reads mapped to TEs in the putative target promoters. The controls (red dots) are the percentages of genome regions covered by TEs in the promoters. **, P-value <0.01, ***, P-value < 0.001, Student’s *t*-test.

Within those reads, more trans ssDRIP-seq reads were enriched in the Alu element than cis ssDRIP-seq reads (*t* test, P < 2.2e-16, Fig 6B). Similar enrichment can also be seen in the trans ssDRIP-seq reads mapped in the 594 predicted target promoters, i.e., on average, about 46.4% (out of 15,592) trans and 33.5% (out of 19,302) cis ssDRIP-seq reads mapped in the Alus (*t* test, p= 1.85e-05, Fig 6C), respectively. In fact, ssDRIP-seq reads were only enriched in the Alu elements in the target promoters, while they were depleted in the TE-free regions (Fig 6D). This is consistent with R-loop formation at Tf2 retrotransposons (Hartono *et al*, 2018) and the cDNA of endogenous TY1 retrotransposons in yeast (El Hage *et al*, 2014). Altogether, our analysis supported the possibility that R-loops at the cAlu elements may be associated with EPIs.

### The cAlu-cAlu pairing in EPI may be subject to strict evolutionary constraint in primates

A strong coevolution signal evidences the functional relevance of cAlu-cAlu pairing. We defined 772, 7977 and 114928 orthologous enhancers (Fig EV3A), promoters and introns, respectively, between human and chimp (see Methods). Based on those orthologs, we further defined 869, 8274 and 221330 conserved Alus in enhancers, promoters and introns, respectively. (Fig EV3B; see Methods).

First, we found that these conserved Alus were enriched in the promoters and enhancers, compared to introns (χ^2^ test, P-value< 2.2e-16; OR>1.24, Fig 7A). The promoter regions were enriched for old Alus. Studies have already shown that the ancient Alu tends to enriched in the enhancers (Su *et al.*, 2014). We then checked the validity of this in promoters. The distribution of Alu subfamilies in enhancers and promoters was remarkably alike (PCC=0.94, p= 5.47e-05, Fig 7B), i.e., the oldest subfamily, Alujo, was enriched for both enhancer and promoters, while the youngest subfamily, AluY, was depleted in those cis-regulators. This analysis suggested a negative selection for recent Alu insertions in both enhancers and promoters.

**Figure7.**
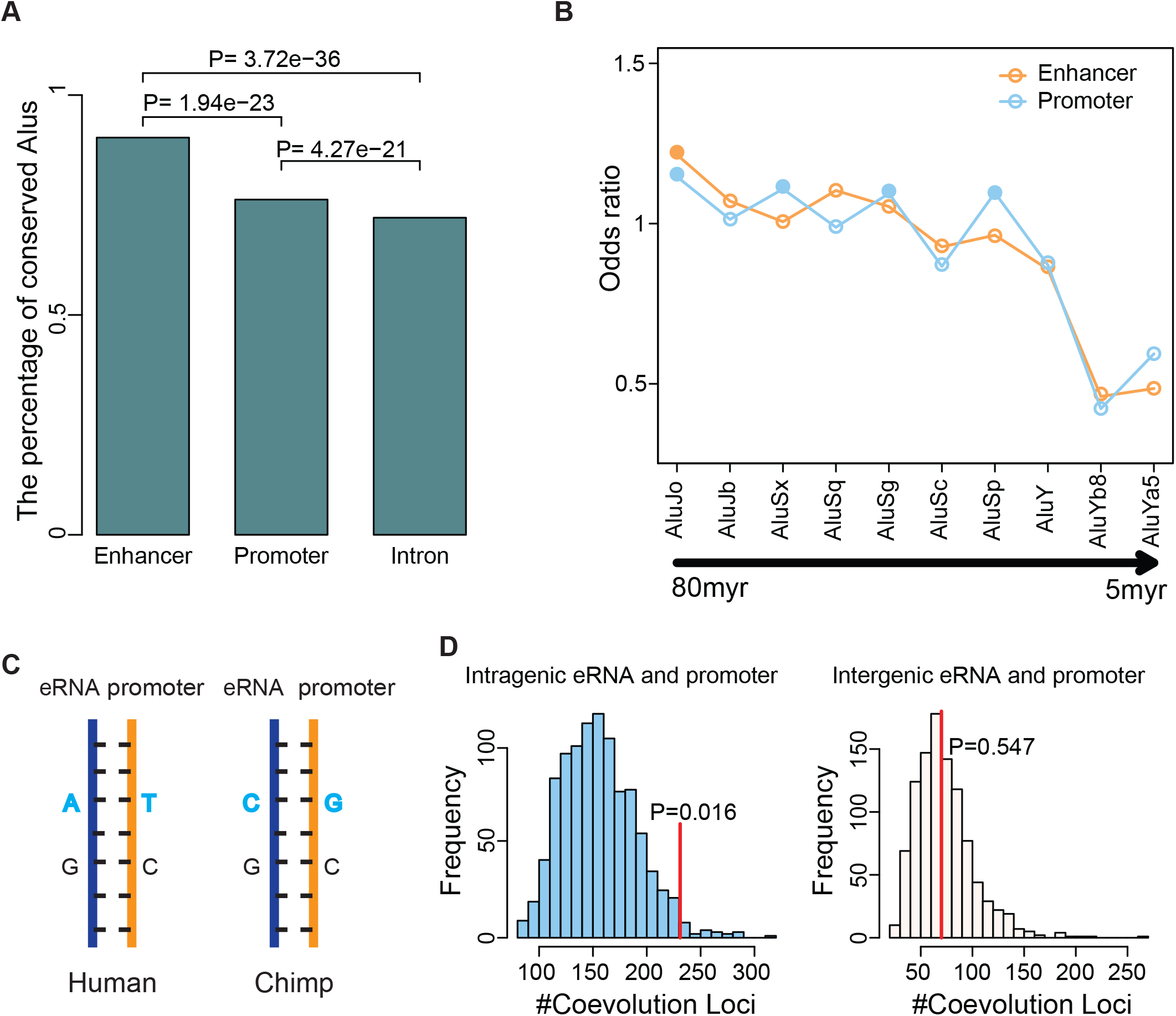
Conservation of cAlu elements. **A.** Percentage of conserved Alus in the orthologous enhancers, promoters and introns between human and chimpanzee (two-tailed χ^2^ test). **B.** Ages of Alu elements in enhancers and promoters. Solid circles indicate significant enrichments (hypergeometric test, P-value<0.01). **C.** The diagram on coevolution. **D.** The enrichment of coevolution loci in EPI. EPIs were grouped into intragenic (left) and intergenic (right) panels. Bars represent the distribution of coevolution event between randomly rewired EPIs, and a total 1000 sets of rewiring were performed in each plot. The red lines indicate observed coevolution events between human and chimpanzee; the z-score equals 2.15 and −0.12 for intragenic and intergenic EPIs, respectively.

Second, enriched coevolution loci were found between cAlu-cAlu pairs. Coevolution is a strong indicator of functional connections, and it has been broadly reported in RNA loops and the interface of protein-protein interactions (Goh & Cohen, 2002; Wen *et al*, 2020). If cAlu-cAlu base-pairing is biologically relevant, evidences of coevolution will be seen between closely related species, e.g., human and chimp (Fig 7C). By comparing the cAlu-cAlu region of orthologous enhancer-promoter interactions between human and chimp (see Methods for details), we identified 231 and 70 coevolution events at the cAlu in the intragenic and intergenic EPIs, respectively (Fig 7D). A significant enrichment for the coevolution event was detected in the intragenic cAlu (P-value=0.016, Fig 7D), compared to permutation control. Considering the fast turnover of cis-regulators during evolution (Domene *et al*, 2013), the orthologous definition of intergenic EPIs in the present work is obviously oversimplified such that little enrichment for coevolution events was found for intergenic cAlus. The indistinguishable coevolution level in the intergenic EPIs may merely suggest the need for a more sophisticated definition for those fast-evolving elements. Nevertheless, our analysis provides strong evidence for the existence of functional base-pairing between eRNA and promoter at the cAlu, at least for the intragenic EPIs.

### Link between aberration at Alu element and disease

If cAlu-cAlu paring is functionally relevant, we would expect the enrichment of disease-causing SNPs in the cAlus. To make this determination, we collected 45288 disease-associated SNPs (pSNP) from the GWAS catalog (MacArthur *et al*, 2017) and 70930 benign SNPs (nSNPs, i.e., combined likely benign and benign SNPs) from the dbSNP database (Smigielski *et al*, 2000), respectively. Indeed, compared to the nSNPs, pSNPs are more enriched in promoter and enhancer Alus, and the odds ratio in promoter is larger than that in introns (promoter, OR=12.47; enhancer, OR=1.76; intron, OR=2.11; introns are from protein-coding genes, and the length of introns is larger than 3kb and less than 50kb; Fisher’s Exact Test, Appendix Table S5-S7).

In particular, we identified SNP site rs9306335 (T/C) in an intergenic enhancer Alu element. This enhancer has four potential target genes, including TAB1, MGAT3, MIEF1 and RPS19BP1. When site rs9306335 was T and C, the four target genes and eRNA were mismatch and perfect match in the BLAST alignment (Fig EV4). SNP rs9306335 was annotated as related to the glycosylation of immunoglobulin in the GWAS catalog. TAB1 is an immune-related gene, and MGAT3 is a glycosylase gene, which plays a crucial role in the glycosylation process of immunoglobulin.

## Discussion

In the present work, we examined multiple models and identified the trans-acting R-loop as the most likely explanation for the dominance of the Alu element in the aligned sequence between enhancers and promoters. Evidences showed that DNA:DNA and RNA:DNA triplex hybrids were depleted from the Alu elements, after controlling for the low mappability of the regions. However, the RNA:DNA hybrids showed clear preference towards the Alus. Moreover, we detected strong signals of coevolution at the aligned Alus between human and primate, implying the functionality of those base-pairings. For the first time, we provided evidences that the sequence per se may play roles in promoter-enhancer interaction, in addition to acting as the binding platform for DNA- and RNA-binding proteins.

We did not predict the exact EPI in this work, we only assessed the statistical characteristics of all intra-TAD enhancer and promoter pairs. This mainly resulted from our limited understanding of active enhancers and their interactions. Although many algorithms have been published in the literature to predict active enhancers, targeted genes, and the interactions between them, the methods remain far from perfect. Furthermore, precise enhancer and EPI prediction may not be that essential in this study because the quality of data from various sources is heterogeneous (Dataset EV1). For example, some data types like GRID-seq were not even generated in the main cell type that we discussed (GM12878), and the RNA:DNA triplex was purely computationally predicted. Thus, to characterize the overall features of EPI is a more practical strategy than one seeking precise prediction. Even so, our data clearly showed the trends, which, in later studies, can be reinforced by more accurate enhancer, eRNA, and targeted promoter predictions.

We noticed that only a small portion of enhancers/eRNAs and EPIs has the features we characterized in this work. As we mainly examined only one cell type (GM12878), which is a lymphoblastoid cell line, we cannot draw the conclusion that the trans-acting R-loop is rare in the other cell types or tissues. It has been reported that the transcription activity of TE is substantially elevated in testis and brain (Pasquesi *et al*, 2020). Therefore, it would be very interesting to explore whether trans-acting R-loops are more prevalent in those tissues. Interestingly, in ssDRIP-seq data on hESC-derived hNSC cells, the author observed enrichment of R-loops on those TEs (Yan *et al*, 2020). On the other hand, R-loops have long been considered a contribution to genome instability (Sollier & Cimprich, 2015). However, increasingly accumulated evidences and examples have shown that R-loops may also function as potential regulators of gene expression (Skourti-Stathaki & Proudfoot, 2014). Thus, more wet experiments will be critical to further establish this trans-acting R-loops model for these parts of EPIs.

Our model does not exclude the utility of existing models for eRNA promoter interactions. In almost all eRNA promoter interaction models, the binding to RBPs is key to the functionality of eRNAs. The R-loop can be bound by the RBPs. In a recently study, Cheung and colleagues identified more than 300 RBPs with preferential binding to R-loops, both *in in vitro* and *in vivo* (Wang *et al*, 2018). Moreover, the combination of RNAs and proteins has been suggested in a recently derived theory (Yamamoto *et al*, 2020) and has, in several cases (Garcia-Jove Navarro *et al*, 2019; Pessina *et al*, 2019), been observed to drive the formation of condensates in the nucleus. Thus, to decipher the function and mechanism of RNA, as well as the hybrids of RNA:DNA, in forming the condensates, would be a rather interesting question. More studies will be performed in the functional genome side of this trans-acting R-loop model of EPIs. We only briefly explored the possible functionality of R-loop-mediated EPI by GO analysis and identified “T cell activate”, “regulation of innate immune response”, “activation of innate immune response” and “leukocyte cell-cell adhesion” as enriched GO terms. N6-methyladenosine (m6A) chemical modification was potentially depleted on the R-loop loci of EPIs in our preliminary exploration (data not shown). It has already been reported that m6A is negatively correlated with A-to-I RNA editing (Xiang *et al*, 2018), which is well known to be enriched in Alus. Thus, it would be interesting to integrate functional genomic data, such as the epitranscriptomics, RNA editing, R-loop and RNA-RBP interaction data, to see how trans-acting R-loops may function in transcription, splicing, and other nuclear processes. Needless to say, having more data generated in more cell types, tissue types and species will be critical to further explore the function and the evolution of those structures.

Taken together, our trans-acting R-loop-mediated EPI model provides a new perspective on the role of eRNAs and Alu elements in the EPI regulation, but more work is needed to fully understand these components in the context of gene regulation networks in mammals.

## Materials and Methods

### Data

The human lymphoblastoid cell line GM12878 was taken as the model system in this study. We summarized the data we used in Dataset EV1.

The NA12878 phased genome was downloaded from http://sv.gersteinlab.org/NA12878_diploid/NA12878_diploid_2012_dec16/NA12878_diploid_genome_2012_dec16.zip.

The SNP locations were downloaded from http://sv.gersteinlab.org/NA12878_diploid/NA12878_diploid_2012_dec16/CEUTrio.HiSeq.WGS.b37.bestPractices.phased.hg19.vcf.gz.

### Identification of eRNA and target genes

The active enhancer regions were defined as the distal overlapping peaks of ChIP-seq of two histone modification marks (H3k27ac, H3k4me1) and DNase I hypersensitive sites (DHS), but with H3K4me3 depleted (Fig EV1C). The peaks not overlapping with promoter regions (TSS-2KB, TSS+2KB) were defined as distal. The ChIP-seq peaks were called by MACS2 with parameters (--nomodel --shift −37 --extsize 73) (Zhang *et al*, 2008), and the DHSs were downloaded from ENCODE (Consortium, 2012). The full-length of enhancer-associated RNAs (eRNAs) was determined as follows. The TSS of eRNA is defined by the Cap Analysis of Gene Expression AND deep Sequencing (CAGE) data (Djebali *et al*, 2012), and the TTS of eRNA is defined as the end of the overlapping region of ChIP-seq peaks. The expression level of eRNA (RPKM) was calculated using the Global nuclear Run-On sequencing (GRO-seq) data (Core *et al*, 2014). The eRNAs with RPKM greater than 0.5 were considered to appear in the sample (Fig EV1A–EV1D), and the transcripts shorter than 200nt, or longer than 2000nt, were excluded from this study. All intra-TAD enhancer-promoter pairs were assumed to be interacting, and the annotation of TADs was downloaded from Rao et al. (Rao *et al.*, 2014) (Fig EV1B, Dataset EV1). The promoters are defined as the [−2k, 1kb] region flanking the TSSs. These strategies have also been applied in K562, HepG2, HeLa-s3 and MCF-7 cell lines.

Since we did not have CAGE data in primate, the definition of enhancer regions in chimp cell line LCL was largely reliant on evolutionary conservation to human. The identified active enhancers in GM12878 were liftOver to chimp genome (pantro5), and their activity was determined using histone marks in chimp with the thresholds identical to those in GM12878. The ChIP-seq and DNase-seq data used for chimp lymphoblastoid cell line LCL were also summarized in Dataset EV1.

### GRID-seq and iMARGI data analysis

The GRID-seq reads were mapped to hg19 with default parameters by STAR (Dobin *et al*, 2013). The reads count for the multi-mapping reads were evenly distributed to each mapped locus. We calculated the accumulated RNA reads numbers per 0.1 kb in the Alu elements of eRNAs and their flanking region. GRID-seq reads were classified into three types, i.e., distal, proximal and interchromosomal ones that the genomic distance between read1 and read2 is larger and smaller than 2kb, respectively, and the inter-chromosomes ones.

The iMARGI peaks were downloaded from the original paper (Yan *et al.*, 2019). The control was selected as the genome fragments in the Alu-free promoters and those having distances to the TSS identical to those of cAlus.

#### Orthologous promoters and enhancers

The orthologous promoters between human and chimp were defined as the [-2k,1k] genome regions flanking the TSSs of the orthologous genes in the public database NCBI HomoloGene (Coordinators, 2017) and could be liftOver from hg19 to pantro5 with at least 90% overlapping in sequence. The orthologous enhancers between Human and chimp were defined as the genome fragments that could be liftOver from human to chimp and were regarded as transcriptionally active, as indicated by histone marks and DNase hypersensitive assay in both species. The base-pairing between enhancers and promoters was determined by BLAST with default setting in each species, independently. The control for enhancer-promoter pairs was generated by randomly shuffling 1000 times.

### Enrichment of CTCF motif orientation

Enrichment of CTCF motif orientation was defined as

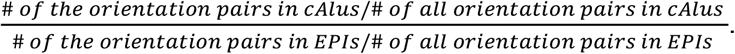

Only 170 EPI cAlu-cAlus contain CTCF motif binding sites. The CTCF motifs were identified by FIMO with default parameters (p-value threshold, 1e-04), and the PWM were downloaded from JASPAR (Fornes *et al.*, 2020).

#### Conserved Alu

Orthologous introns were defined as the regions that were liftOvered from the introns in hg19 and overlapped with at least 90% of the annotated Chimp intron sequence in Pantro5 in the orthologous genes, as defined in HomoloGene (Coordinators, 2017).

#### The conserved cAlus were defined as

Alus in orthologous promoters, introns or enhancers.

a. Alus in Human and chimp belong to the same Alu subfamily.
b. In the case of multiple Alus in promoter/enhancer, the rank and distance to TSS of the conserved cAlus will be similar between Human and chimp (Fig EV3A).

### CTCF ChIP-seq mapping

CTCF ChIP-seq data were mapped by Bowtie2 (version 2.3.4.1) with default parameters, except −k 10. Reads to be counted in each locus have been mapped. Reads with low mapping quality and duplicate reads were removed by samtools (version 1.9). Peaks were called by MACS2 (version 2.1.1.20160309) with default parameters.

### TAD

The annotation of TADs for GM12878, K562 and HeLa cells was downloaded from GSE63525 (Rao *et al.*, 2014); for MCF-7 and HepG2 cells, the TADs were called by Juicer (Durand *et al*, 2016) (Dataset EV1). The TADs for chimp were liftOvered from the TAD annotation in GM12878 to Pantro5.

### SNP data

GWAS SNPs (release v1.0.3) were downloaded from the GWAS catalog(MacArthur *et al.*, 2017). The disease-associated SNPs were manually curated by reading the annotation for traits. Benign SNPs were downloaded from dbSNP (Build 151) (Smigielski *et al.*, 2000). We manually curated benign SNPs by choosing the entries with annotation “Benign” and “Likely benign” in the Clinical Significance. All SNPs can be found in Dataset EV4.

## Acknowledgements

We are grateful to Dr. Yang Yu, Dr. Jing Li, Dr. Junfeng Liu, Dr. Bingxiang Xu, Dr. Hui Zhang, Yiqun Zhang and Xiao Zhang for helpful discussions. Mr. David Martin performed English language editorial services. This work was supported by the National Nature Science Foundation of China (91940304, 31871331, 31671342), Beijing Natural Science Foundation (Z200021), Special Investigation on Science and Technology Basic Resources of MOST, China (2019FY100102), the National Key R&D Program of China (2018YFC2000400), and the Beijing Advanced Discipline Fund (115200S001).

## Author contributions

XB and ZZ conceived this project. XB developed the method and analyzed data, XB and ZZ prepared the manuscript. All authors read and approved the final manuscript.

## Data availability

This study includes no data deposited in external repositories. The analysis code involved in this work can be found at https://github.com/BaixueXK/workspace.

**Figure EV1. Identification of eRNAs and their target genes.**

**A.** Workflow chart for eRNA identification.

**B.** All intra-TAD promoter-enhancer interactions were considered in this study.

**C.** Ratio of H3k4me3 to H3k4me1 reads in the overlapped peaks. The red bars were the peaks with enriched H3k4me1 and depleted H3k4me3.

**D.** An example of eRNA. The peaks of histone marks were called by MACS2 and marked as black horizontal lines in the corresponding rows. We marked three CAGE peaks as A, B and C that located in the overlapping region of H3k4me1 and H3k27ac peaks. A was used to define the TSS of an eRNA in the plus strand, and B and C were used to define the TSSs of two eRNAs in the minus strand. The 3’end of eRNAs was defined as the boundary of the overlapping regions of H3k4me1 and H3k27ac peaks. Putative eRNAs were drawn in the last row.

**E.** Significantly more EPIs were supported by ChIA-PET loops. In the same TAD, the regions with the same length as that of enhancers were randomly selected to form control EPI with the target promoters (U test).

**F.** The accumulated contact frequencies of predicted EPI and controls. The controls were taken from random pairing intra-TAD bins with identical distribution of genomic distance to the EPIs (two-tailed two-sample *t*-test).

**Figure EV2. Gene expression level in different cell lines.**

**A-D.** Cumulative frequency of putative target gene expression for K562, HepG2, HeLa-s3 and MCF-7 cells, respectively. E+ and E-represent the enhancers carrying at least one or zero Alu element, respectively. P+ and P-represent the putative target promoters carrying at least one or zero Alu element, respectively. The blue dotted line represents the EPIs in the E-P+ group after removal of genes with multiple Alu-carrying enhancers.

**Figure EV3. Orthologous enhancer and conserved Alu between human and chimp.**

**A.** Definition of orthologous enhancers between human and chimp.

**B.** Definition of conserved Alu and non-conserved Alu. Alus were regarded as orthologous when they belong to the same Alu subfamily, have similar ranks in the orthologous anchors, and have similar distance to the start or end locus of the anchor. The color represents an Alu subfamily. The start and end of anchors were marked by the black stars and triangles, respectively. The anchors could be either orthologous enhancers, promoters or introns.

**Figure EV4. Example of a GWAS SNP located at intergenic eRNA and related to IgG glycosylation.** Reference allele is T, and the mutation allele is C. The target gene TAB1 is an immune-related gene, and MGAT3 is a glycosylase gene.

